# The impact of a psychedelic drug on olfactory search behavior by mice

**DOI:** 10.1101/2025.07.09.663970

**Authors:** Amanda C. Welch, Nate Gonzales-Hess, Takisha A. Tarvin, Jessica Connor, Matthew C. Smear

## Abstract

Animals use their sense of smell for survival and well-being. Any disruption to olfactory perception, such as in the case of olfactory hallucinations, can lead to devastating consequences and decreased quality of life. Psychedelics interrupt and distort accurate perception, yet little is known about the impact of psychedelics on olfactory behaviors. Using an olfactory search task, we investigated the impact of the psychedelic 2,5-dimethoxy-4-iodoamphetamine (DOI) on olfactory search behavior of mice. We found that DOI decreases search accuracy, alters movement, and increases sniff rate. These findings suggest that the olfactory behaviors are altered by DOI, elucidating psychedelic-induced changes to olfactory processes.

## Introduction

Self-movement induces changes in perception. Animals control and exploit these perceptual changes by active sensing (Ahissar & Assa, 2016; Buzsáki, 2019; Nelson & MacIver, 2006). Hallucination is another form of perceptual change. In a hallucination, the ability to differentiate between reality or hallucination may depend on active sensing (J. J. Gibson, 1962; James J. Gibson, 1970). Capturing active sensing as a behavioral output can give insight into internal states under the influence of psychedelics.

Olfactory disturbances are well documented in numerous psychiatric and neurological disorders, including schizophrenia, Parkinson’s disease, depression, epilepsy, and migraines (Bannier et al., 2012; Kopala et al., 1994; Mainardi et al., 2017; Sarnat & Flores-Sarnat, 2016). Patients with these diagnoses may experience olfactory hallucinations, where prevalence varies, with some reports as high as 34.6% (Kopala et al., 1994) and a sex difference showing women more frequently affected than men (Bainbridge & Byrd-Clark, 2020; Sjölund et al., 2017). Olfactory hallucinations are predominantly unpleasant, with patients most often reporting smoke, waste, decaying matter, and mold (Sjölund et al., 2017). Their largely negative impact motivates research to explore olfactory hallucinations. While hallucinations occur in psychiatric and neurological disorders, they may also be induced by the consumption of psychedelics. Here, we introduce a mouse model of behavior for exploring psychedelics and their impact on olfaction.

Psychedelics induce stereotypical behaviors in both humans and animals. In rodents, administration of a serotonergic psychedelic reliably produces a rotational head shake known as the head-twitch response (HTR; (González-Maeso et al., 2007; Halberstadt et al., 2020; Halberstadt & Geyer, 2013). The quantity of HTRs correlates with the potency of hallucinations in humans. This correlation has motivated the idea that HTRs are a behavioral analog to the human experience (Hanks & González-Maeso, 2013). In humans, erratic, high-rate eye saccades occur after administration of a classic psychedelic (Hebbard & Fischer, 1966). In schizophrenia, auditory hallucinations can be predicted by subtle lip movement (Rapin et al., 2013), and humming can decrease the rate of auditory hallucinations (M. F. Green & Kinsbourne, 1990).

Considering the importance of olfaction in rodents and the large gap in understanding how psychedelics impact the olfactory system, we sought to establish an understanding of respiration and behavior of mice after the administration of the psychedelic DOI (2,5-Dimethoxy-4-iodoamphetamine). Here, we show the changes in active sampling and olfactory search behavior after the administration of the psychedelic DOI. Taken together, our results lay the foundation for using mouse olfaction in behavioral preparations for understanding the mechanisms of psychedelics.

## Results

### DOI alters olfactory search behavior

To study the impact of psychedelics on active olfactory behavior we administered DOI to mice and tested them in an olfactory search paradigm in which they report the source location of a noisy, turbulent odor gradient (Findley et al., 2021). In the search paradigm, mice poked into an initiation port to start a trial. Odor was pseudo-randomly delivered from the left or right side of the opposite end of the arena. To receive a water reward, mice had to navigate toward higher odor concentration. To investigate task performance in the absence of a stimulus, we omitted odor on 10% of trials. Mice were trained until they performed at 85% accuracy for 1 week, which varied per mouse but took less than 2 weeks total. Mice were then administered 3 mg/kg DOI or saline then placed into the task 5 min later (Figure 1A). Mice were crossed over into the opposite condition 1 week later.

**Figure 1.**
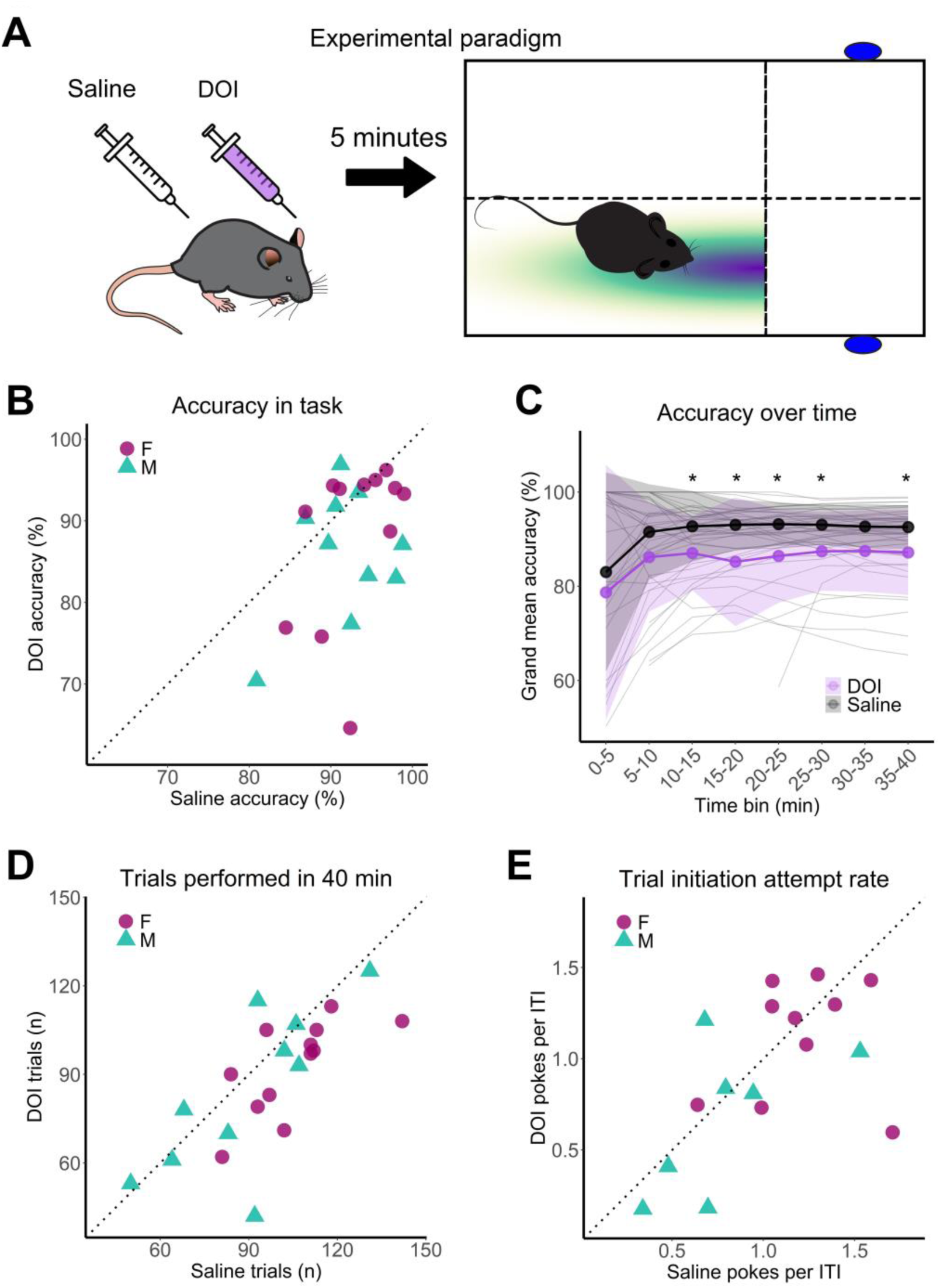
Mice performance in the olfactory search paradigm. **(A)** Diagram of experimental paradigm. Mice were injected with 3 mg/kg DOI or saline then placed into the task 5 min later. **(B)** Scatter plot of accuracy in the task for each mouse in DOI and saline conditions. Color and shape denote mouse sex. **(C)** Line graph of grand mean accuracy in the task binned in 5 min intervals. Shaded areas are standard deviation. **(D)** Scatter plot of number of trials performed for each mouse. Sessions were at a fixed duration of 40 min. **(E)** Scatter plot of trial initiation attempts during the ITI for each mouse. Rate was normalized as the number of pokes into the initiation port by the number of ITIs.

Given that psychedelics disrupt perception in humans, we hypothesized that mice would perform less accurately in olfactory search. Indeed, mice more often navigated to the incorrect side of the arena in the DOI condition (Figure 1B; Wilcoxon signed rank test, p = 0.024, *n* = 22). DOI was administered 5 min before the session began, so we wondered whether its effect would become larger as the session proceeded. To ask how DOI impacted performance as a function of time in the session, we quantified accuracy in 5 min time bins. In odor trials, accuracy was higher in the saline condition after the first 10 min, except for the 30 – 35 min bin (Figure 1C; Wilcoxon rank-sum test, p < 0.01 for time points within 10 – 25 min, p = 0.039 for 35 – 40 min). Thus, the change in performance is overall uniformly distributed across the duration of behavioral sessions.

Since high doses of DOI can decrease locomotor activity in an open field test (Halberstadt et al., 2009), we next assessed task adherence after DOI administration. Mice completed fewer trials in the DOI condition (Figure 1D; Wilcoxon signed rank test, p = 0.011, *n* = 22). This effect was driven by female mice; female mice performed fewer trials on DOI compared to saline (Figure 1D; Wilcoxon signed rank test, p = 0.011, *n* = 12), whereas male mice did not show a significant difference (Figure 1D; Wilcoxon signed rank test, p = 0.39, *n* = 10). We further assessed whether mice repeatedly poked into the initiation port during the ITI to start a new trial. We found that mice poked into the initiation port, but there was not a significant difference between conditions (Figure 1E; Wilcoxon signed rank test, p = 0.27, *n* = 17), indicating an equivalent level of motivation to perform the task.

In an auditory detection task, mice false alarmed at a higher rate both when a signal was expected and when administered the dissociative psychedelic drug ketamine compared to vehicle (Schmack et al., 2021). Similarly, we hypothesized that mice in the DOI condition would report an odor stimulus during the ITI of a session, especially during a longer ITI, but this was not the case. Mice rarely entered the reward port during the ITI after trial initiation attempt (Wilcoxon rank sum test, p = 1). These results demonstrate that mice have an impaired ability to accurately complete an olfactory search task, but persist in trial initiation and completion, indicating a psychedelic-induced disruption to olfactory-guided behavior.

### DOI impacts speed and timing

When given the drug ketamine to induce hallucination-like percepts, mice wait longer for a reward when they are confident they experienced a stimulus, even when one did not occur (Schmack et al., 2021). This may be due to confidence as a function of time invested awaiting a variably scheduled reward. Since mice cannot explicitly report confidence in a percept, this metric allows for a measurable output that they were confident in their decision. If psychedelics impact confidence in our olfactory search task, we would predict that the mice would spend longer in the reward port in the DOI condition. Reward port duration was increased in the DOI odor trial condition compared to both saline odor and saline odor omission trials (Figure 2A; Wilcoxon rank sum test, p < 0.001, *n* = 17). We did not find a sex effect of condition (Figure 2A supplemental, p = 1). When comparing the median duration between conditions, mice in the DOI condition spent longer in the reward port compared to saline (Figure 2B; Wilcoxon signed rank test, p = 0.002). This effect was driven by the odor trial (Figure 2B supplemental; Wilcoxon signed rank test, p = 0.001), as we did not see a difference between median duration after an omission trial (Wilcoxon signed rank test, p = 0.60).

**Figure 2.**
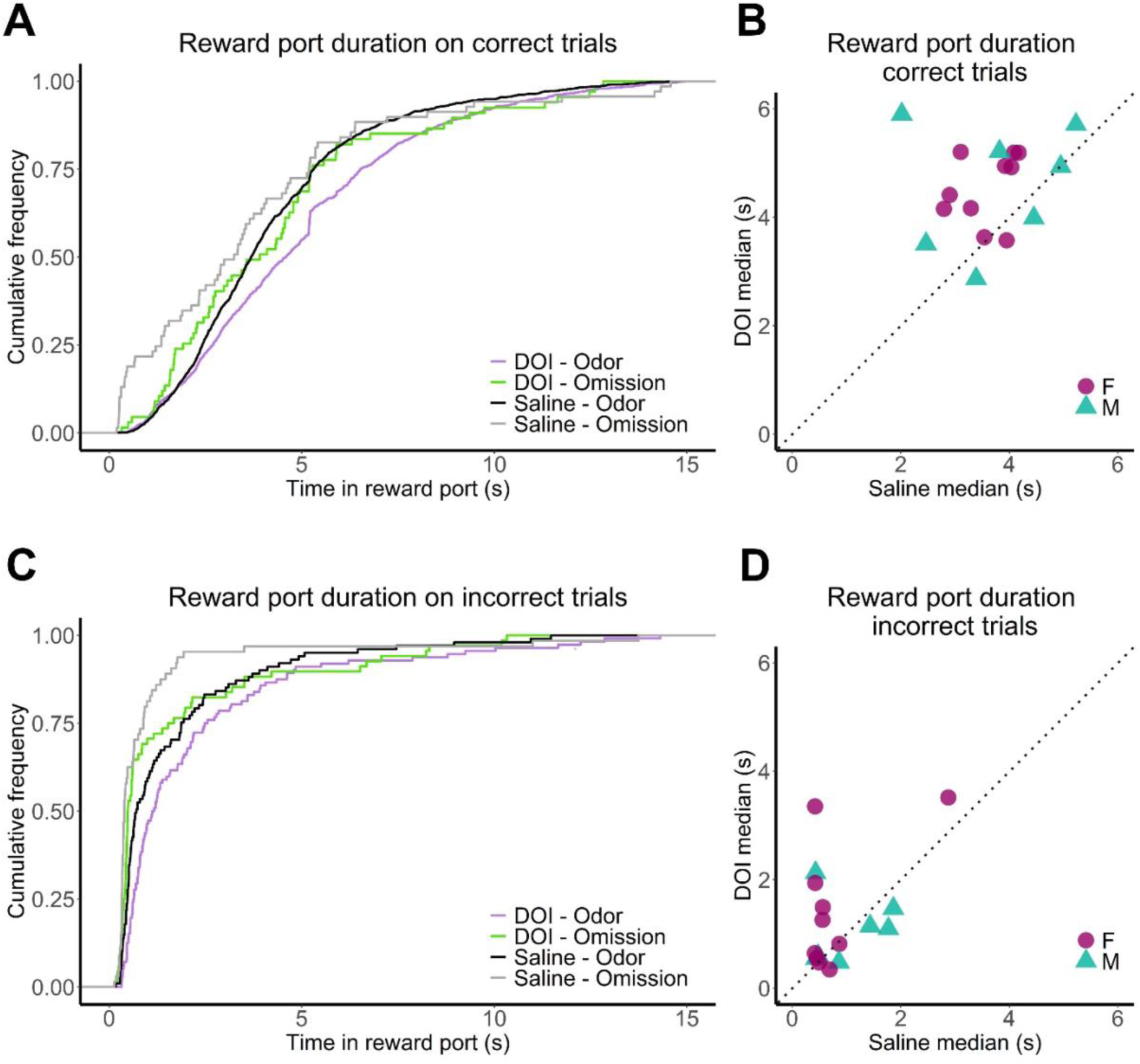
Duration in reward ports increased in the DOI condition. **(A)** Cumulative histogram of duration in a reward port after a correct response in a trial. Colored lines show DOI odor and omission trials. Black and grey lines show saline odor and omission trials. **(B)** Scatter plot of median duration in a reward port after a correct trial per mouse in DOI and saline conditions. **(C)** Cumulative histogram of duration in a reward port after an incorrect response in a trial. Color as in A. **(D)** Scatter plot of median drinking duration after an incorrect trial per mouse in DOI and saline conditions.

In incorrectly performed trials, when mice did not receive a reward, mice spent the longest duration in the reward port after the DOI odor trial and the least amount of time in the reward port after a saline odor omission trial. We found an increased duration in DOI odor trials compared to DOI odor omission trials (Figure 2C; Wilcoxon rank sum test for all comparisons, p < 0.001), saline odor trials (p = 0.030), and saline odor omission trials (p < 0.001). Additionally, there was an increased reward port duration for the saline odor trial compared to the saline odor omission trial (p < 0.001). Sex differences within each condition were not significant (Figure 2C supplemental, Wilcoxon rank sum, p = 1). When comparing the median duration between conditions, there was no difference between DOI and saline (Figure 2D; Wilcoxon signed rank test, p = 0.21). However, after an odor omission trial, mice spent longer in the reward port in the DOI condition (Figure 2D supplemental, Wilcoxon signed rank test, p = 0.049). Taken together, we show that mice spend longer in the reward port under the influence of DOI. Our results are thus consistent with previous work suggesting that psychedelics may increase perceptual confidence (Schmack et al., 2021). However, we cannot rule out a general impact of psychedelics on task adherence. An increased reward port duration could also be due to reduced vigor in movements out of the reward port. DOI can increase dopamine signaling and reward salience (Martin et al., 2024), thus the increased duration in the reward port may be driven by an increased valuation of water.

Since mice show decreased locomotion at high doses of DOI (10 mg/kg) (Halberstadt & Geyer, 2018) we asked whether trial duration and movement speed varied between conditions. In odor trials, mice took longer to complete a trial in the DOI condition compared to the saline condition (Figure 3A; Wilcoxon signed rank test, p < 0.001, *n* = 22). In contrast, there was no difference in trial duration in omission trials between DOI and saline (Wilcoxon signed rank test, p = 0.26, *n* = 22). Next, we wanted to know if mice took longer to initiate a new trial and thereby increased time in the ITI. However, there was no difference in ITI duration between conditions (Figure 3B; Wilcoxon signed rank test, p = 0.19, *n* = 22).

**Figure 3.**
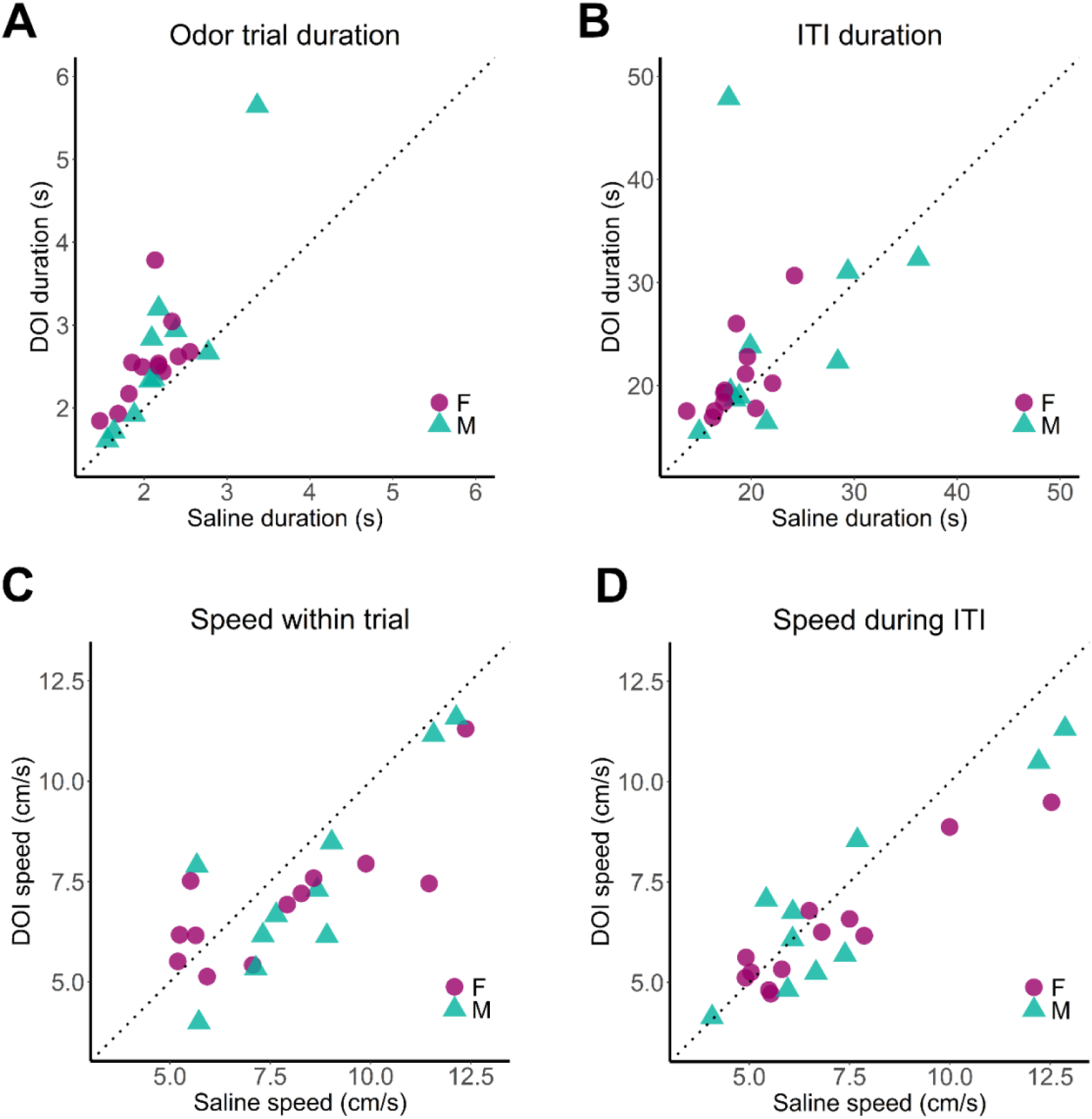
Trial duration increased and speed decreased in the DOI condition. **(A)** Scatter plot of mean odor trial duration for each mouse in the DOI and saline condition. **(B)** Scatter plot of mean ITI duration. **(C)** Scatter plot of mean speed within trial for each mouse in the DOI and saline condition. **(D)** Scatter plot of mean speed during the ITI.

Using DeepLabCut (DLC; (Mathis et al., 2018), the nose, head, and center of mass were tracked from video. We found that DOI decreased movement speed both within trial (Figure 3C; Wilcoxon rank sum test, p = 0.011, pooled trial type) and during the ITI (Figure 3D; Wilcoxon rank sum test, p = 0.036) compared to saline. In odor trials specifically, the speed between DOI and saline sessions were not different (Figure 3 supplemental; Wilcoxon rank sum test, p = 0.059). Within the DOI condition, speed was decreased in the ITI compared to an odor trial (Figure 3 supplemental; Wilcoxon signed rank test, p < 0.001). However, there was no difference in speed between the odor and odor omission trials in the DOI condition (Wilcoxon rank sum test, p = 0.21), nor the ITI and odor omission trials (Wilcoxon rank sum, p = 0.068). In comparison, in saline sessions the ITI speed was slower compared to odor (p = 0.002) and odor omission trials (p = 0.017). Speed within the odor and odor omission trials were not different in the saline condition (Wilcoxon rank sum, p = 0.54). Taken together, the increased duration within the reward ports, the increased trial duration and decreased speed demonstrate a change in interaction with the task, indicating that psychedelics may influence olfactory perception, confidence, or movement vigor.

### DOI increases sniff frequency

Active sensing is crucial to investigating the environment. Rodents increase sniff rate when investigating a novel odor or in an olfactory task (Findley et al., 2021; Kepecs et al., 2007; Verhagen et al., 2007; Youngentob et al., 1987). In humans, administration of LSD increases eye movements (Hebbard & Fischer, 1966), but does not alter respiration rate (Sokoloff et al., 1957). We next assessed the impact of DOI on sniff rates. When all sessions with sniff measurement were combined, sniff frequency was higher in DOI sessions (Figure 4A – C; Fisher-Pitman Permutation test, Z = 24.9, p < 0.001). All mice with a matched DOI and saline session had a higher-shifted distribution sniff rates in the DOI condition (Figure 4 supplemental; Kolmogorov-Smirnov test, p < 0.001 for 13 of 13 mice). When comparing sniff frequency by trial type, mice sniffed faster in the DOI condition for each trial type and during the ITI (Figure 4D; Fisher-Pitman Permutation test for all trial types; Odor trials: Z = 3.38, p < 0.001; Omission trials: Z = 2.71, p= 0.007; ITI: Z = 18.4, p < 0.001). We did not observe a difference in median sniff frequencies between conditions (Figure 4E; Wilcoxon signed rank test; p = 0.11). Since mice rely on sniffing to investigate and determine the source of an odor stimulus, and mice increase sniff rate during an investigation phase of search, the sustained increase in sniff rate suggests that a psychedelic drug impacts olfactory perception.

**Figure 4.**
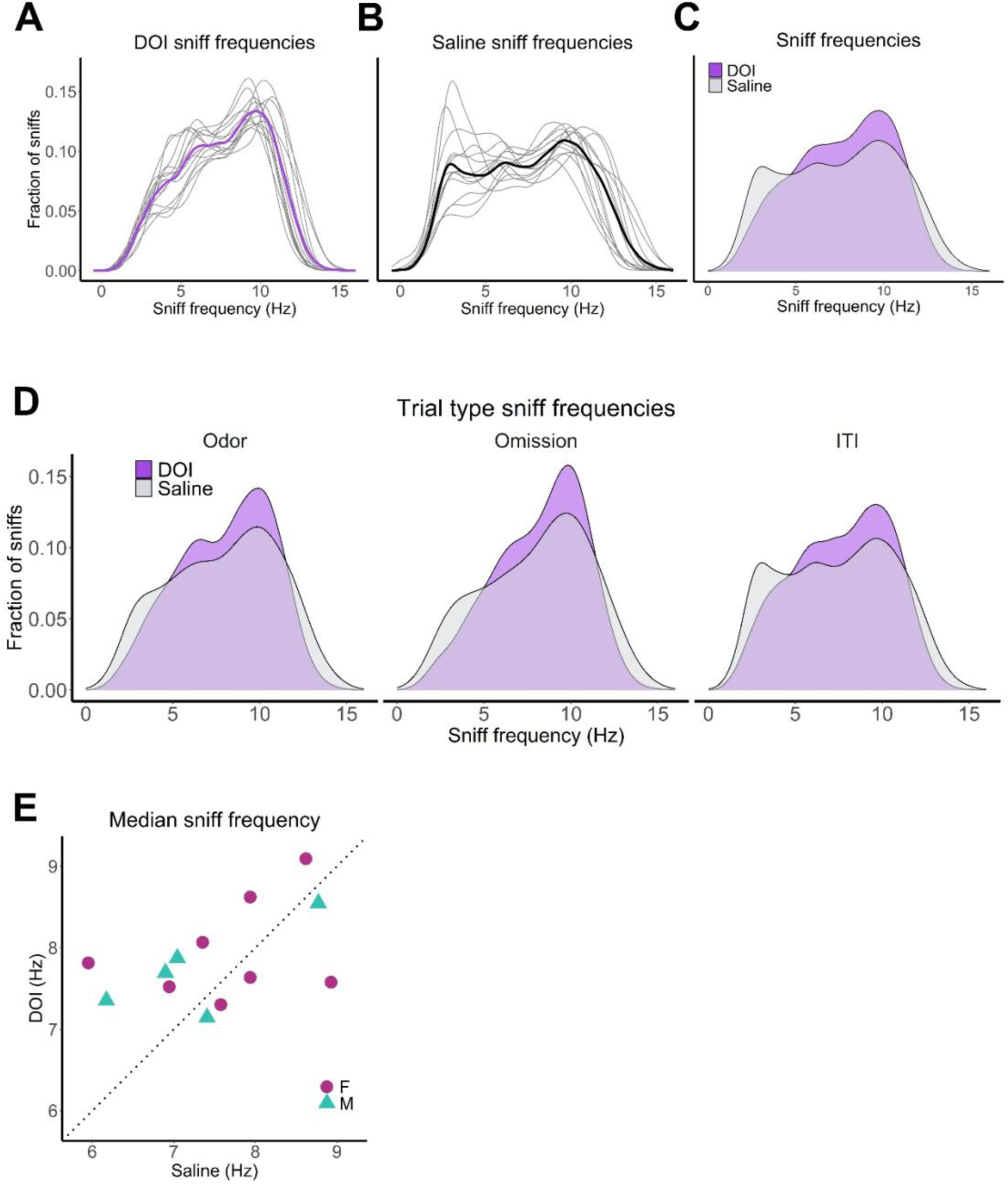
Sniffing frequency increased in the DOI condition. **(A)** Histogram of sniff frequencies across all mice with signal in the DOI condition (*n* = 14). The thick purple line represents all mice, thin grey lines are individual mice. **(B)** Histogram of sniff frequencies across all mice with signal in the saline condition (*n =* 13). The thick black line represents all mice, thin grey lines are individual mice. **(C)** Histogram of all DOI and saline sniff frequencies overlaid. Purple is the DOI session, grey is the saline session. **(D)** Histograms of sniff frequencies for an odor trial, odor omission trial, and during the ITI. Purple is DOI, grey is saline. **(E)** Scatter plot of median sniff frequency per mouse in the DOI and saline condition.

### Mice occupy allocentric space similarly but head twitch preferentially in proximity to odor ports

To investigate the allocentric structure of behavior under these conditions, we quantified occupancy across all mice. We hypothesized that DOI would alter task adherence, which predicts that mice would spend more time in areas irrelevant to the task such as corners of the arena, with a more even distribution across the arena. Contrary to this prediction, mice showed similar occupancy across DOI and saline conditions (Figure 5A, B).

**Figure 5.**
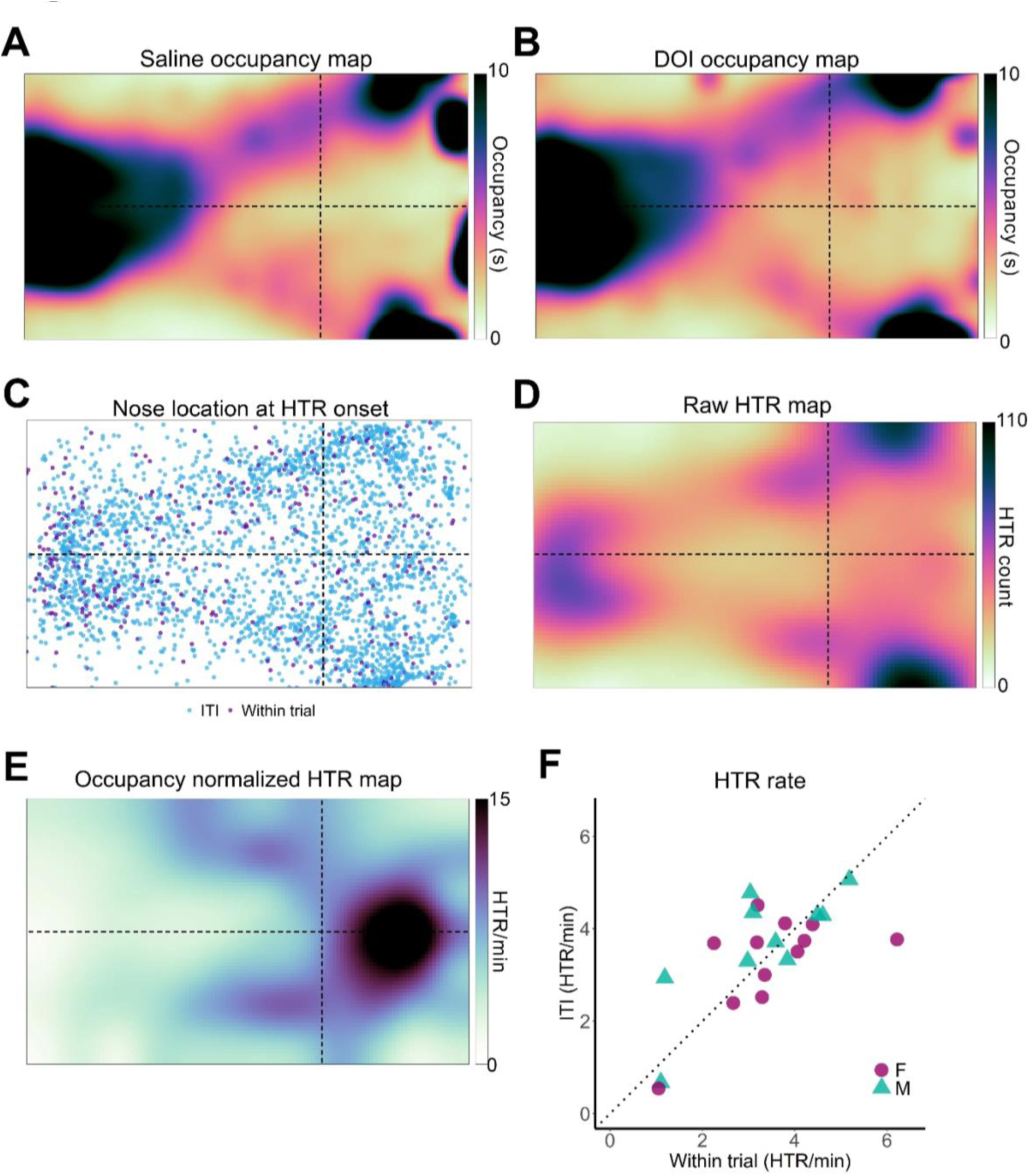
Mice similarly occupied the arena between conditions, but head twitched more frequently near odor ports. **(A)** Two-dimensional histogram of occupancy (fraction of frames spent in each 1 cm^2^ bin) in the saline session. Color is mean across all mice (*n* = 22). **(B)** Same as in A, but for the DOI session. **(C)** Scatter plot of nose location at the onset of a HTR across all mice. **(D)** Two-dimensional histogram of HTR location and Gaussian smoothed. **(E)** Two-dimensional histogram of HTR rate during a DOI session based on ∼2.5 cm^2^ bin location. HTR was normalized based on occupancy per mouse, then averaged across all mice (*n =* 22). Color bar represents the HTR rate per min. **(F)** Scatter plot of HTR rate per mouse in the ITI versus within a trial.

The head-twitch response (HTR) is a reliably observed behavior when mice are administered a psychedelic. Next, we assessed where in the arena mice emitted HTRs. If HTRs are merely an ectopic movement independent of sensory perception, we would predict that their rate of emission should be uniform throughout the arena. If HTRs depend on stimuli and behavior, we would predict that their probability would vary in different positions. We plotted the nose location at the onset of a HTR (Figure 5C) then spatially binned the arena in ∼ 2.5 cm^2^ to quantify HTR location counts (Figure 5D). We normalized the rate of HTR based on occupancy per mouse, as some areas of the arena had disproportionate levels of occupancy. During a trial, HTR rate occurred throughout the arena without spatial predictability on the Y-coordinate (Figure 5 supplemental; Permutation test, p = 0.92), but with preference for the X-coordinate (Figure 5 supplemental; Permutation test, p = 0.002). Similarly, during the ITI, we found that mice HTR more frequently closer to the end of the arena after crossing the decision line (as X-coordinate increases), but not immediately next to the reward ports (on the Y-coordinate) (Figure 5 supplemental) and when conditions were combined (Figure 5E; Permutation test, p < 0.001 for X-coordinates). Hallucinations that occur in Parkinson’s disease have impairment of the Dorsal Attention Network and Default Mode Network coupled with sensory regions (Shine et al., 2015), suggesting hallucinations occur more due to attention disruption. Thus, in the search phase of a task, we expected HTR frequency to be lower. We normalized HTR rate based on within trial and ITI duration and found no difference in HTR rate based on whether the mouse was in a trial or ITI (Figure 5F; Wilcoxon signed rank, p = 1). Additionally, we did not see a sex difference in HTR rate over the 40 m session (Figure 5 supplemental; Wilcoxon rank sum, p = 0.31). Taken together, these results indicate that the HTR is not a randomly occurring movement tic, but is tied to a high odor probability location in the arena.

## Discussion

In this study, we demonstrate that the psychedelic DOI impacts olfactory navigation and breathing rhythms in mice. Mice continue to perform the olfactory search task, but their performance statistics demonstrate a decreased ability to correctly report odor source location. Female mice, but not male mice, performed fewer trials in the DOI session than in the saline session. Timing in the task is skewed slower in the DOI condition. Mice move more slowly and take more time to complete an odor trial in the DOI condition, as demonstrated by trial duration. Mice also collect rewards differently as demonstrated by time in the reward port. On incorrect trials when there is no water reward delivered, mice still spend a significant amount of time in the reward port after a DOI odor trial. These findings give insight into how a psychedelic drug impacts olfaction.

Several timing metrics are altered in mice on DOI, suggesting that DOI may change their sense of time. Psychedelics interfere with the perception of time in mice (Halberstadt et al., 2016) and reduce reaction time in humans (Yousefi et al., 2025). Additionally, mice that are confident they experienced a stimulus wait longer in a reward port for a reward (Schmack et al., 2021). The increased duration in the reward port on incorrect odor trials could be a sign of increased confidence in an incorrect decision. Alternatively, mice may lose track of how much time has been spent in the reward port. The salience of a reward may also be increased during the DOI condition, as DOI increases dopamine signaling for predictable rewards (Martin et al., 2024). Thus, the increased duration in the reward ports could be an increase of the value of water during the task.

Mice rely on sniffing to investigate their environment. In this study, we demonstrate that the sniff rate is altered with DOI in a way that suggests increased active sensing to discern the odor landscape. Sniffing is highly synchronized to movement and investigation (Findley et al., 2021), however, in the DOI condition we see that mice sniff faster throughout the session, despite moving more slowly in the task. This increase in sniff rate is evident in both trial conditions and during the ITI. The increase in sniff rate taken with the decrease in movement speed suggests that sniffing the environment in the absence of an external stimulus could be due to a hallucination-like perception. Mice investigate and sniff novel and attractive odors at a higher rate than neutral or familiar odors (Li et al., 2023; Qiu et al., 2021; Wesson et al., 2008), yet we see a sustained elevated sniff rate in the absence of novel or attractive odors.

The HTR is a reliable behavior observed in rodent psychedelic research. Serotonergic psychedelics induce a HTR rate that varies with dose (Canal & Morgan, 2012). In this study, we expected to observe the HTR at a higher rate during the ITI without location dependency. However, we found that the HTR is location dependent after occupancy normalization but not trial dependent. Hallucinations are argued as a top-down process, where prior assumptions and internal state can influence perception (Grossberg, 2000; Kowalski et al., 2024; Powers et al., 2016). The proximity to the location of the odor source may thus contribute to the increased HTR rate. As mice approach the decision line in a trial, odor concentration in the air becomes higher, suggesting the prior knowledge of an odor source could influence perception. However, there is uncertainty in what brain regions are directly responsible for the HTR. The HTR can be induced by infusion of a psychedelic into the prefrontal cortex (Willins & Meltzer, 1997), while transection of the brain at the posterior commissure eliminates the HTR (Bedard & Pycock, 1977). Whether the HTR is purely a motor tic or a behavior in response to perceptual experience is unknown. However, if the HTR were happening randomly as a tic might do, we would not expect to see spatial preference for this behavior.

Overall, this work advances our understanding of how the psychedelic DOI alters olfactory search behavior and sniff in freely moving mice. Our findings demonstrate that psychedelics impact olfactory behavior and thus establish a framework for studying hallucination-like percepts in olfaction. This work sets the stage for future investigation of how psychedelics influence neural dynamics in the olfactory system.

## Materials and methods

### Animal housing and care

All experimental procedures were approved by the Institutional Animal Care and Use Committee (IACUC) at the University of Oregon and in accordance with the ethical guidelines of the National Institutes of Health. Animals were group housed on a reverse 12:12 h light cycle. All behavioral testing was performed during the dark cycle. Food and water were given ad libitum unless otherwise specified for water restriction. Mice were given a minimum of 1 mL water daily and maintained above 80% pre-water restriction weight. Health was monitored daily and removed from water restriction if weight dropped below 80% baseline as approved by the IACUC. All mice used were C57Bl6/J and at least 12 weeks of age, but no more than 5 months, at the time of surgery.

### Surgical procedure

Animals were deeply anesthetized with isoflurane (3% concentration during induction, then maintained at 1-2%) for the procedure. The incision site was topically anesthetized with lidocaine (8 mg/kg) SQ prior to incision. Meloxicam-ER (4 mg/kg) and buprenorphine (0.1 mg/kg) were given SQ peri-operatively. Supplemental lactated ringers (LRS, 1 mL) were given SQ post-operatively to improve recovery and surgical outcome. Health was monitored daily for 3 days post-operatively.

Thermistors were implanted between the nasal bone and inner nasal epithelium to measure respiration (Findley et al., 2021). A custom titanium head bar was secured to the skull. Prior to surgery, the thermistor (TE Connectivity, #GAG22K7MCD419) wires were minimally stripped then soldered into pins (JST Sales America, # A02KR02DS28W305B) for signal conduction. The wire was fixed in place using cyanoacrylate. Exposed tissue was sealed using Vetbond tissue adhesive. Exposed skull was sealed using cyanoacrylate. A small dot of orange tempera paint (Pro Art, #4435-2) was stamped onto the headbar for live tracking in Bonsai.

### Drugs

Drugs were administered IP 5 min prior to behavioral testing. (±)-DOI hydrochloride (DOI) (Sigma-Aldrich, #D101) was dissolved in sterile physiological saline and injected at a fixed volume of 5 mL/kg at a dose of 3 mg/kg. Prior to administration, freshly mixed drugs were sterilized through a 0.2-micron filter. Sterile physiological saline was used as the control (Hospira, #NDC 0409-4888-02).

### Behavioral testing

For full documentation on rig software and use protocols, see: https://nghess.github.io/fmon-docs/

### Water association

Mice were water restricted and trained to associate 3 pokes in the behavioral arena with water. Each poke was calibrated daily to dispense 6-8 µL water, each within 0.1 µL of the other. Water was made available in an alternating fashion: initiation, left, initiation, right, repeat. Each session lasted 30 min. Mice were moved to the odor navigation task after completing 50 pokes.

### Odor navigation task

Mice were water restricted and maintained at a minimum of 80% baseline weight for training and experiments. They were trained first to associate an initiation port, left reward port, and right reward port in an alternating pattern with a water reward. When mice received a minimum of 50 water rewards in a 30-min session by poking into these ports, they graduated onto the odor navigation task. In the two-alternative choice odor navigation task, no water was awarded by initiating a trial, only when correctly navigating to the left or right side of the arena where the odor was produced. A correct response was registered when the mouse entered the corresponding quadrant where odor was released. An overhead camera tracked movement in real-time and provided live feedback to the system. Upon correctly navigating to the left or right, mice had 5 seconds to poke into the reward port to receive their reward. The left and right sides were calibrated each day to ensure equivalent volumes for each reward, ranging from 1.2 to 1.8 µL. We pseudo-randomly assigned left or right-side trials with a maximum of 5 consecutive trials on one side. The odorant 2-Phenylethanol (2-PE) (Sigma-Aldrich, #77861) was diluted in deionized water to a final concentration of 0.001%. Control trials without odor administration occurred at a rate of 10% total trials. We also imposed a variable inter-trial interval (ITI) of 1 to 15 s regardless of correct response. Mice were able to complete as many trials as possible within 40 min. When mice were able to complete 5 consecutive sessions with an accuracy of 85% or higher to ensure task comprehension, mice were assigned to start in the DOI or saline condition then crossed over into the other condition at least 1 week later.

### Pose estimation and behavioral syllables

The location of the nose, head, left and right ears, body, left and right hips, and tail base were tracked via DeepLabCut (Mathis et al., 2018). A limit was placed on tracking data so the likelihood of < 0.95 and points that moved faster than 8 pix/frame were smoothed by linear interpolation. Movement speed was calculated as the distance traveled (difference in position) over time elapsed (s). Tracked videos were analyzed in Keypoint Motion Sequencing (KP-MoSeq; (Wiltschko et al., 2020) to obtain behavioral syllables.

### Head-twitch response

Raw video of behaving mice was played at 50% speed in Adobe Premiere Pro. The HTR was manually marked by an observer when at least 3 paroxysmal rotations of the head occurred. The onset of the HTR (+/- 2 frames) was when the time stamp was marked. Time stamps were exported to csv for alignment with tracked pose estimations and further analysis.

### Data analysis

Analysis of sniff signals were cleaned in MATLAB and visualized in R. Inhalation and exhalation times were extracted by finding peaks and troughs in the temperature signal after smoothing with a 25 ms moving window. Any sniffs with dropped out exhalation times were excluded. Analyses of DLC tracking, performance statistics, and sniff frequencies were performed in R. KP-Moseq analyses were performed in MATLAB.

### Figure 1

Accuracy (percent correct) was calculated by dividing the correct odor trials by total odor trials in both the DOI and saline conditions. Accuracy for odor omission trials was calculated with the same method. Rolling accuracy was calculated by dividing the current number of correct trials by the current iteration of trials for each trial for each mouse. Sessions were separated into sequential 5 min time bins. The mean accuracy for trials within each time bin was calculated for each mouse in the odor, odor omission, DOI, and saline conditions. The grand mean was calculated taking the mean of all the means from each mouse at each time bin. Trials completed were calculated by taking the sum of all trials in a session per mouse. Repokes were calculated by taking the sum of initiation port beam breaks during the ITI. A false odor report was calculated as the occurrence of an initiation port beam break followed by crossing the decision line during the ITI. The number of mice used in repoke and false report analyses was 17 due to data acquisition updates that were not in place for the first cohort of mice.

Statistical tests were performed using R Statistical Software (v 4.4.1). Graphs were made using package ggplot2 (Wilkinson, 2011). The Wilcoxon signed rank test was used for paired samples and group comparisons. The Wilcoxon rank-sum test was used for rolling accuracy data since some mice did not perform during a time bin, creating unmatched data. Paradigm figure was created using images from https://scidraw.io/ and https://stock.adobe.com using Adobe Photoshop.

### Figure 2

We used 17 mice in our drinking duration calculation due to our initial system not capturing beam breaks in the reward ports. All mice after our first cohort of 5 had reward port breaks recorded. For correct trials, reward port duration was calculated as time poking into the reward port until exiting the reward port. Re-entries into the reward port after exit did not add time to the calculation. For incorrect trials, duration in the reward port was calculated with the same method although no water reward was delivered.

Statistical tests were performed in R. For the reward port duration cumulative frequencies, normality was checked using the Shapiro-Wilk test. One or more groups violated normality assumptions. The Kruskal-Wallis was used to test for a difference in drinking duration groups. Post-hoc pairwise comparisons were performed using the Wilcoxon rank sum test with Bonferroni adjustment for multiple comparisons to reduce Type I error. Wilcoxon signed rank tests were used for paired comparison of median durations.

### Figure 3

Trial duration was calculated by subtracting trial start time from trial end time for DOI and saline conditions. Trials were grouped by odor or odor omission trials. The mean duration was taken for trial type and condition for each mouse. Actual ITI duration was calculated as the difference between the current trial start time and previous trial end time. The mean ITI duration was taken for each condition and mouse. Task speed was determined using DLC keypoints. Tracking data was imported and points with likelihood < 0.95 or delta > 8 pixels were interpolated. Interpolation and speed calculations were used with modified functions from the collection DLC Analyzer (Sturman et al., 2020). Any speed that was > 160 cm/s was excluded from analysis. Using the nose keypoint, mean speed for each condition per mouse was calculated by taking the distance traveled over time elapsed.

Statistical tests were performed in R. For trial durations, ITI duration, and overall task speed, Wilcoxon signed rank tests were used for paired comparison.

### Figure 4

Initial pre-processing of sniff was performed in MATLAB. Inhalation and exhalation times were extracted by finding peaks and troughs in the temperature signal after smoothing with a 25 ms moving window. Sniffs with duration less than the 5^th^ percentile and greater than the 95^th^ percentile were excluded from analysis. Exported sniff signals with inhalation and exhalation time stamps in ms were imported into R using the R.matlab package (Bengtsson, 2022). Sniff duration was calculated by taking the difference between inhalation start times of successive sniffs. Any sniff event with missing inhalation or exhalation time stamps were excluded from analysis. Any sniffs below 10 s or above 1000 s were excluded from analysis. Sniff frequency was calculated as 1000/difference in consecutive inhalation time stamps. Trial summary files that stored trial start and trial end time stamps were imported into R. Sniff inhalation and exhalation times were converted to sec and categorized as within trial or ITI sniffs based on trial start and end time stamps.

Not all sessions recorded had usable sniff signals due to variable levels of noise, signal drop out, or signal loss. For mice with a usable sniff recording in both DOI and saline sessions, sniff frequencies were compared between conditions within subject using the Kolmogorov-Smirnov test in R. Since some mice had usable sniff in one condition but not the other, we ran a two-sample Fisher-Pitman Permutation test with 10,000 permutations on all usable sniff frequencies between DOI and saline conditions. We ran the same test for all trial types: odor trial, omission trial, and ITI. Permutation tests were performed using the coin package in R (Hothorn et al., 2008).

### Figure 5

Occupancy was calculated by binning the arena into 1 cm^2^ bins, then mean duration in each bin for each mouse. Grand mean was calculated for occupancy in both conditions. HTR rate was calculated by binning the arena into 2.5 cm^2^ bins. HTR was normalized to occupancy by counting HTR occurrences in each bin, then diving that number by occupancy for each mouse. The final normalized HTR map was calculated as the grand mean of normalized occupancy. Plots were upscaled then Gaussian smoothed in MATLAB and colored using the Cubehelix package (D. A. Green, 2011). Statistical testing was performed in R. Coordinate bin labels were permuted 10,000 times and HTR rate compared between the original observed values and permuted values.

## Acknowledgements

We would like to thank members of the Smear Lab for their contributions to this work.

---

**Figure 2 supplemental.**
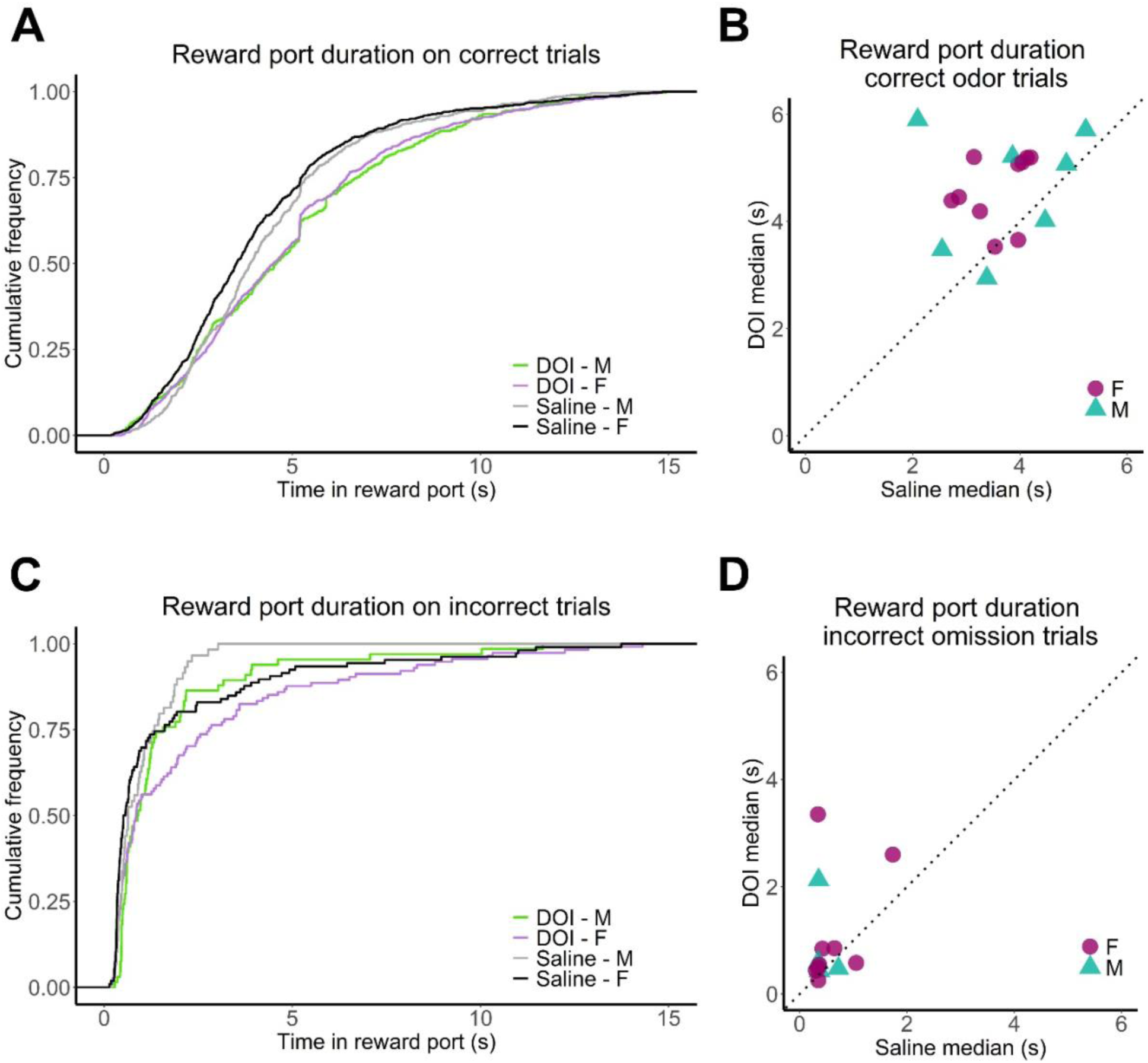
Reward port duration was not sex dependent. **(A)** Cumulative histogram of duration in a reward port after a correct response in a trial. Colored lines show DOI trials separated by mouse sex. Black and grey lines show saline trials by sex. **(B)** Scatter plot of median duration in a reward port after a correct odor trial per mouse in DOI and saline conditions. **(C)** Cumulative histogram of duration in a reward port after an incorrect response in a trial. Color as in A. **(D)** Scatter plot of median drinking duration after an incorrect odor omission trial per mouse in DOI and saline conditions.

**Figure 3 supplemental.**
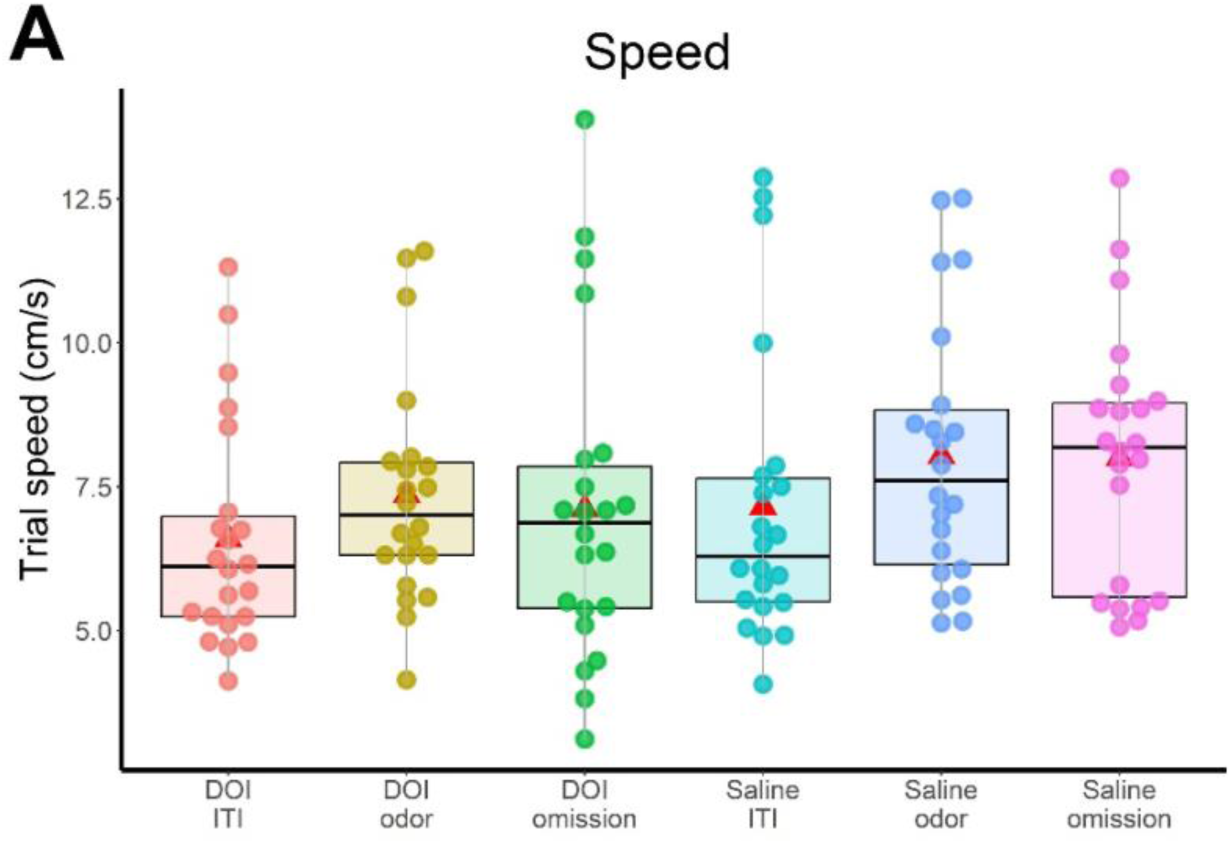
Speed across all conditions and trials. Whisker plots of speed for each mouse in each drug condition and trial type.

**Figure 4 supplemental.**
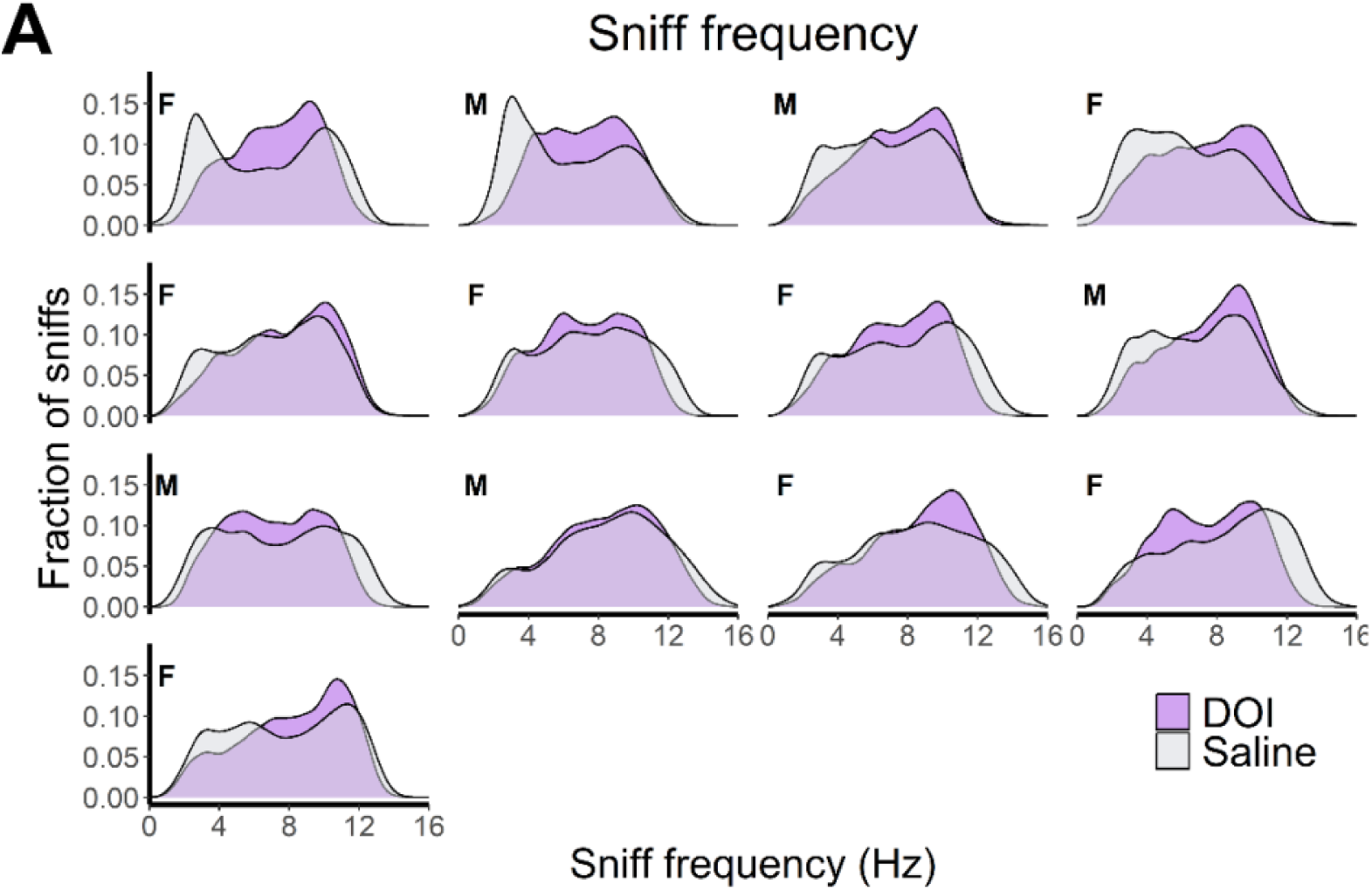
Sniff frequencies per mouse. Histograms of sniff frequency for each mouse with a DOI and saline condition.

**Figure 5 supplemental.**
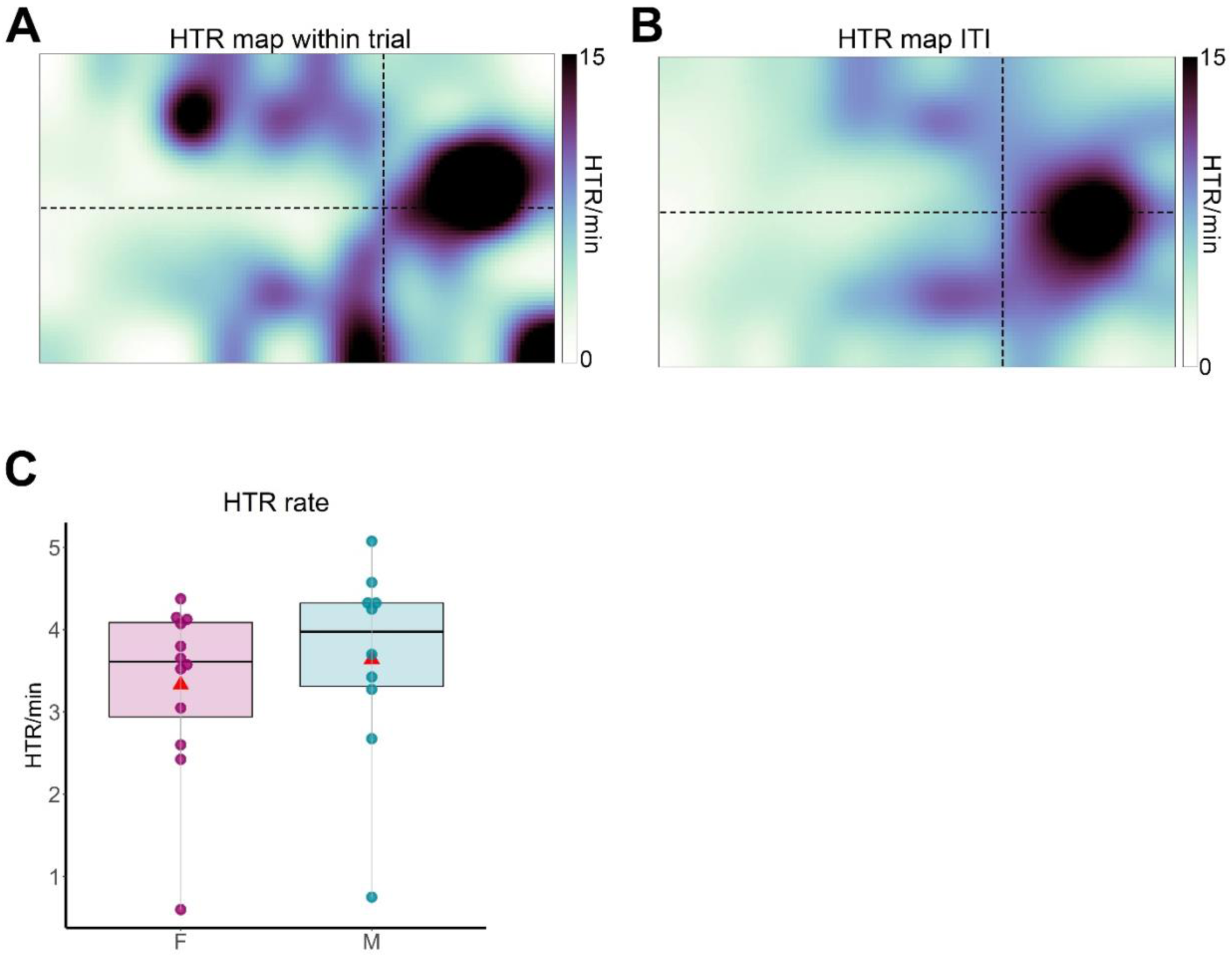
Head-twitch response during the trial and inter-trial interval. (A) Two-dimensional histogram of HTR rate within a trial in the DOI session based on ∼2.5 cm^2^ bin location. HTR was normalized based on occupancy per mouse, then averaged across all mice (*n =* 22). Colorbar represents the HTR rate per min. **(B)** Same as in A, but for the ITI. **(C)** Whisker plot of HTR rate between male and female mice. Red square represents the mean rate.

